# Defining the anti-Shine-Dalgarno sequence interaction and quantifying its functional role in regulating translation efficiency

**DOI:** 10.1101/038984

**Authors:** Adam J. Hockenberry, Adam R. Pah, Michael C. Jewett, Luís A. Nunes Amaral

## Abstract

**Summary:** Studies dating back to the 1970s established that binding between the anti-Shine-Dalgarno (aSD) sequence on prokaryotic ribosomes and mRNA helps to facilitate translation initiation. The location of aSD binding relative to the start codon, the full extents of the aSD sequence, and the functional form of the relationship between aSD binding and translation efficiency are important parameters that remain ill defined in the literature. Here, we leverage genome-wide estimates of translation efficiency to determine these parameters and show that anti-Shine-Dalgarno sequence binding increases the translation of endogenous mRNAs on the order of 50%. Our findings highlight the non-linearity of this relationship, showing that translation efficiency is maximized for sequences with intermediate aSD binding strengths. These mechanistic insights are highly robust; we find nearly identical results in ribosome profiling datasets from 3 highly diverged bacteria, as well as independent genome-scale estimates and controlled experimental data using recombinant GFP expression.

## INTRODUCTION

The abundance of different protein species within a single cell can vary by several orders of magnitude, and multiple points of control are critical for tuning the expression of individual proteins over such a wide-range(Dekel and Alon, 2005; Kudla et al., 2009; Salis et al., 2009; Taniguchi et al., 2010). Transcription of the gene of interest is a necessary first step in the pathway of gene expression but, by itself, transcription is insufficient to ensure protein expression; studies in a variety of organisms have shown that mRNA abundances only modestly predict protein abundances (Guimaraes et al., 2014; Lu et al., 2007; Schwanhäusser et al., 2011; Taniguchi et al., 2010; Vogel and Marcotte, 2012; Vogel et al., 2010). Although the magnitude of these correlations remains open to debate, it is clear that the rate at which different mRNA species are translated into their protein product is a significant source of variation in protein abundance and a point of regulation(Csárdi et al., 2015; J. J. Li et al., 2014).

In studies dating back to the 1970s, researchers noted that a strong interaction between the 16S ribosomal-RNA and the 5’ untranslated region (UTR) of mRNAs is important for overall translation efficiency—defined here as the number of protein molecules made per mRNA per unit time—by enhancing translation initiation in prokaryotes(Shine and Dalgarno, 1974). The strength, optimal distance to the start codon, and structural accessibility of this anti-Shine-Dalgarno::Shine-Dalgarno (aSD::SD) interaction all play a crucial role in modulating the rates of translation initiation and thus protein abundance variation(Barrick et al., 1994; Chen et al., 1994; de Smit and van Duin, 1994, 1990; Rinke-Appel et al., 1994). More recently, multiple studies have reinforced this paradigm and continue to elucidate the finer details about the importance of translation initiation signals, such as highlighting the fact that the surrounding nucleotides may constrain SD sequence evolution due to mRNA structural constraints(Barendt et al., 2013; Bentele et al., 2013; Espah Borujeni et al., 2014; Goodman et al., 2013; Hockenberry et al., 2014; Kosuri et al., 2013; Mutalik et al., 2013).

The aSD motif is highly conserved across prokaryotic taxa, indicating a strong conservation of its mechanism(Lim et al., 2012; Nakagawa et al., 2010). However, not all genes within a given species initiate via the aSD::SD mechanism and there is a large heterogeneity of SD sequence usage between different organisms: studies have reported that over 90% of the genes from some organisms are preceded by an identifiable SD sequence while for other organisms this number can be as low as 10%(Calogero et al., 1988; Cortes et al., 2013; Kramer et al., 2014; Nakagawa et al., 2010).

Over the past 20 years, a number of different studies have analyzed various facets of translation initiation sequence variation across bacteria, but definitions about which genes to consider as “SD genes” varies broadly (Chang et al., 2006; Ma et al., 2002; Na et al., 2010; Nakagawa et al., 2010; Sakai et al., 2001; Salis et al., 2009; Starmer et al., 2006; Zheng et al., 2011). The main differences concern where to look upstream of the start codon for a putative SD sequence and what bases of the 16S rRNA sequence to consider as the aSD sequence when assessing binding strength to the 5’ UTR of mRNAs (Figure 1A). Experimental studies present a well-controlled system to interrogate these mechanisms, but it's unclear whether conclusions from experimental investigations of recombinant genes are valid or relevant at the genome-scale given the complexity of interactions and constraints on coding sequence evolution. Towards this end, recent studies have suggested that the aSD binding strength shows no relationship with the measured translation efficiency of endogenous genes at the genome-scale (G.-W. Li et al., 2014; Li, 2015; Schrader et al., 2014).

**Figure 1:**
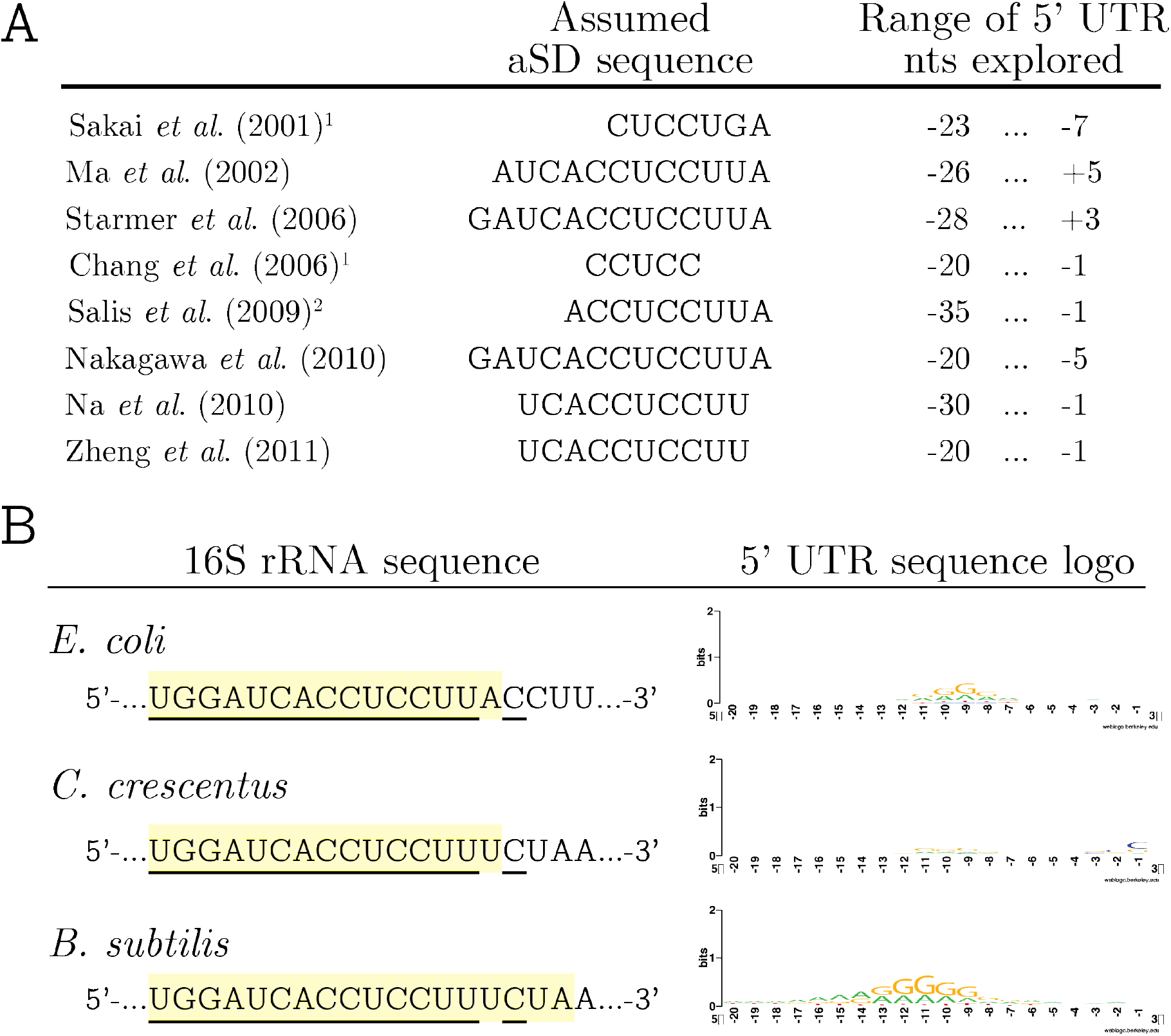
SD sequence usage is variably defined in the literature and differs between genomes. A) Example of several studies that report a range of relevant parameters used to identify the aSD::SD sequence interaction. (^1^denotes studies that implicitly derive aSD sequences by extrapolating from over-represented UTR motifs; ^2^denotes studies that explicitly penalize for non-optimal distances to the start codon). B) The anti-Shine-Dalgarno sequence region in the three organisms studied is highly conserved (left). Underlined bases show perfect conservation across the three organisms; the shaded region denotes the annotated 3’ end of rRNA genes. By contrast, 5’ untranslated regions (UTRs) are largely heterogeneous within and between species (right).

Ribosome profiling enables examination of protein translation for thousands of endogenous transcripts in a single experiment, providing unprecedented insight into the mechanisms of translation at the genome scale (Ingolia et al., 2009). Application of this technique to multiple organisms has already enhanced our understanding of translational regulation, stoichiometric protein production, determinants of elongation speed, and genome annotation(Ingolia et al., 2009; G.-W. Li et al., 2014; Li et al., 2012; Schrader et al., 2014). Here, we investigate whether these ribosome profiling based estimates of translation efficiencies can be leveraged to precisely define the relevant parameters associated with aSD::SD sequence interaction in endogenous genes. Our analysis yields data-driven definitions for the optimal distance of aSD binding to the start codon and the extent of the aSD sequence. We further highlight a highly conserved non-linear relationship between aSD binding and translation efficiency of endogenous genes whereby intermediate aSD binding strengths maximize translation efficiency. We confirm these findings in independent genome-scale and experimental datasets, and in doing so highlight the robustness of our conclusions while validating that the size of this effect is greatly enhanced given less noisy experimental data.

## RESULTS

### Deriving translation efficiency measurements from Ribo- and RNA-seq

For a given mRNA, ribosome density maps derived from ribosome profiling (Ribo-seq) can be used to illustrate regions of relatively fast and slow translation. When used in conjunction with RNA sequencing (RNA-seq) to estimate mRNA abundances, this data can also be used to roughly quantify relative translation efficiency (RTE) on a per gene basis. However, it is important to note that estimates of RNA abundances and ribosome occupancies are both error-prone due to biological noise as well as the numerous steps in the experimental process that may introduce systemic bias(Lahens et al., 2014; Miettinen and Bjorklund, 2014; Mohammad et al., 2016; Steijger et al., 2013; Zupanic et al., 2014). Thus, RTE is a particularly error-prone measurement because uncertainty is compounded when dividing two noisy values. We therefore established several quality controls for gene inclusion (see Materials and Methods). Following on the previous work of others(G.-W. Li et al., 2014; Schrader et al., 2014), we then we calculated relative translation efficiency (RTE) per gene as:

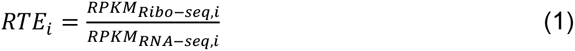

where RPKM_Ribo-seq_ and RPKM_RNA-seq_ are reads per kilobase per million mapped reads for a gene, *i*, obtained through ribosome profiling and RNA-seq, respectively. Using the original Ribo- and RNA-seq mappings provided by three separate studies in rich media for *E. coli, C. crescentus*, and *B. subtilis* we derived measurements of translation efficiency for 2910, 1833, and 2385 genes, respectively (Supporting Fig. 1)(G.-W. Li et al., 2014; Schrader et al., 2014; Subramaniam et al., 2013). While this metric relies on some crucial assumptions, such as equivalent elongation rates between genes, prior work has shown that these assumptions are generally valid(G.-W. Li et al., 2014); a noise-free RTE metric calculated in this manner should be highly correlated with ‘true’ translation efficiencies as we have defined it for all but the most extreme cases of genes with highly irregular elongation and/or termination patterns.

As others have noted, mRNA structure surrounding the start codon is known to influence translation initiation, perhaps playing a dominant role in determining translation efficiency(de Smit and van Duin, 1990; Gu et al., 2010; Hockenberry et al., 2014; Kudla et al., 2009; G.-W. Li et al., 2014). We confirmed this finding by showing that log-transformed translation efficiencies in all three organisms had highly significant correlations with the predicted degree of mRNA secondary structure (Δ*G_folding_*) in the initiation region (defined here as -30 to +30 nucleotides relative to the start codon) (R^2^ = 0.13, 0.10, and 0.08 for *E. coli, C. crescentus*, and *B. subtilis, p*<10^−43^ for all casesj. Given the strength of this correlation (Supporting Fig. 2), moving forward we analyze the residuals from this predictive model (in units of log-scaled translation efficiency) in order to determine what role, if any, aSD-binding strength has in modulating translation efficiency:

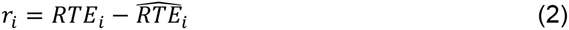

where *RTE_i_* is the relative translation efficiency of gene *i*, and 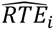 is the estimate for gene_*i*_ derived from the regression on Δ*G_folding_* for each organism. We include this step to alleviate the source of biological variation associated with cis-structure, but note that these computational predictions introduce further technical error due to the at best modest correlation between computationally predicted structures and their *in vivo* counterparts.

### Defining the optimal distance to the start codon and species specific aSD sequences

Using the residual RTE values above, we took a systematic approach in order to determine first where to look, in an unbiased manner, relative to the start codon for the signal of aSD binding under the assumption that the true value of this parameter should show the strongest correlation between binding strength and residual RTE values. For each gene, we calculated the hybridization energy of the core aSD sequence (5’-CCUCC-3’) to each sequential 5-nucleotide segment upstream of the start codon (Fig. 2A). Hereafter, we will refer directly to the location (relative to the start codon) as the number of bases between the farthest 5’ base included in the putative aSD sequence, corresponding to the aligned spacing(Chen et al., 1994). We asked how well the binding energies at a particular location for *all* genes performed at predicting our model residuals via both 1^st^ and 3^rd^ order polynomial regression (to account for a potentially nonmonotonic relationship).

**Figure 2:**
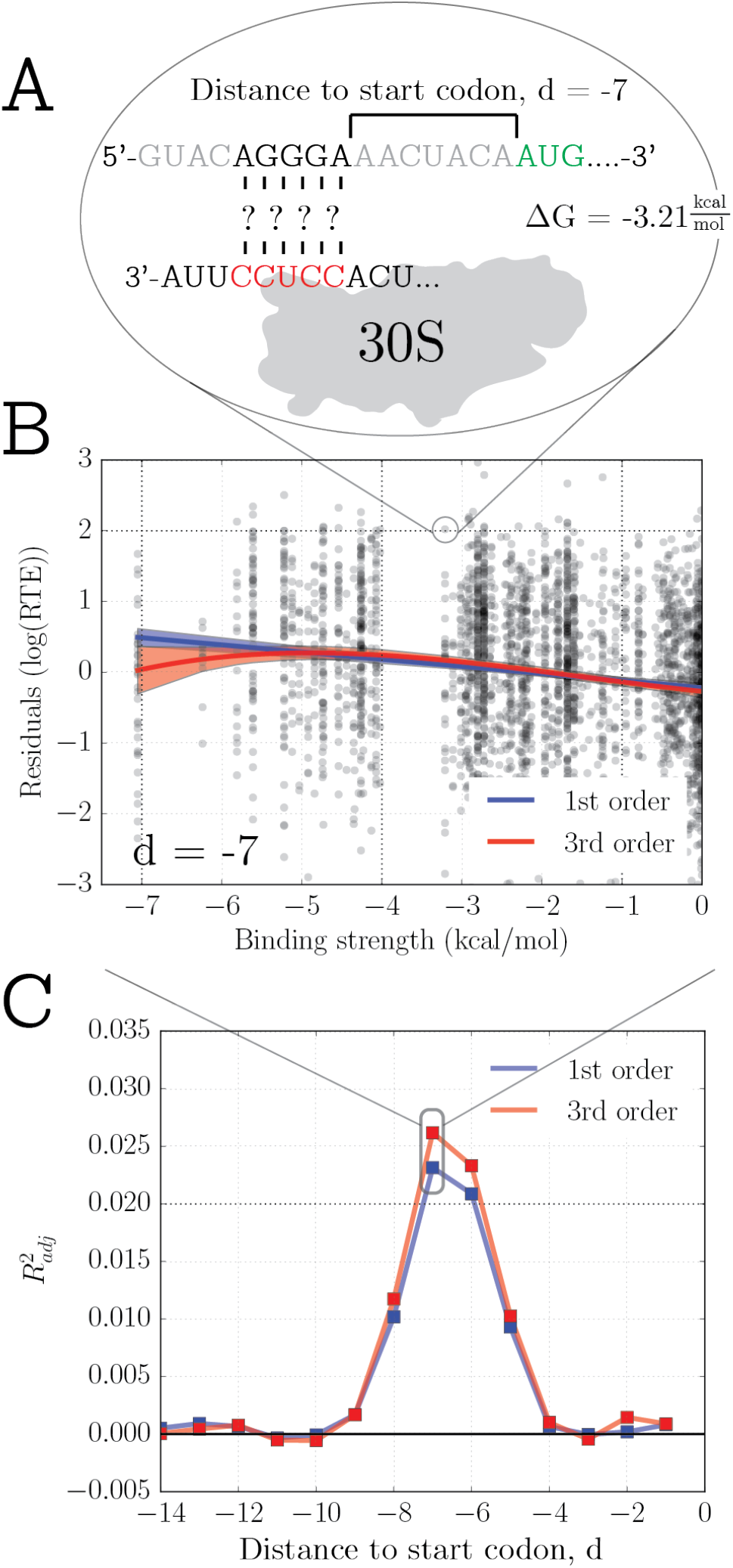
Determining the optimal aligned distance to the start codon using regression analysis on residual RTE values. A) Illustration of the method used in this study for determining the Gibbs free-energy of the hybridization of the putative aSD sequence (highlighted in red) to the 5-nucleotide sequence at a distance of 7 nucleotides upstream from the start codon, indexed from the start codon to the closest base in the hybridization. B) The strength of aSD binding for each gene at a distance of -7 is correlated against the model residuals in units of log(RTE). Shown are 1^st^ and 3^rd^ order polynomials (R^2^_adj_ = 0.023 and 0.026 for the 1st and 3^rd^ order fits, p<10^-16^ for both). C) We performed the same correlation analysis as in (B) for each distance to the start codon in the *E. coli* dataset. Shown are the R^2^_adj_ values for the relevant models with a clear maximum peak for d=−7.

In Fig 2B we show the data for a distance to the start codon of -7 nucleotides (representing 7 unpaired nucleotides before the 5’-aSD base of ‘C’ in this example). We show both the first and third order fits for the translation efficiency data from *E. coli;* both correlations are small yet nevertheless highly significant (F test, *p*<10^−16^). In Fig. 2C, we show the adjusted-R^2^ (R^2^_adj_) resulting from repeating the correlations shown in Fig. 2B for each indicated spacing. We utilize the R^2^_adj_ metric hereafter because unlike R^2^ this adjusted metric penalizes for increasing parameter numbers associated with more complex models and thus helps guard against over-fitting our model to the data. The sharpness of this peak indicates that even despite the measurement noise we are able to detect a clear and highly robust relationship between aSD binding to the 5’ UTR of mRNAs and translation efficiency. The 3^rd^ order polynomial model was slightly more predictive at this stage, so we present our data in the form of 3^rd^ order polynomial regressions hereafter except where otherwise noted.

We repeated the above analysis for different putative aSD sequences extending in the 5’ and 3’ directions at different binding locations and observed significantly enhanced R^2^_adj_ values and a slight repositioning of the optimal distance to the start codon (Fig. 3A,B). We finally explored a range of variants that include extensions on both ends to determine the optimally predictive aSD sequence and distance parameters for the given dataset (Fig. 3B). Several of these putative aSD sequences produced similar results so we used the shortest sequence amongst these candidates (5’-ACCUCCUUA-3’) but stress that our methodology can likely not discriminate these boundaries precisely giving the small differences between putative aSDs with single base additions/deletions. While the overall correlation coefficient in this best fit model is still modest (R^2^_adj_ = 0.041), the significance of this finding is extremely high (*p*<10^−27^) indicating that despite the potentially large error in RTE estimates, we are nevertheless able to observe a highly significant underlying relationship. We further observed that the 3^rd^ order model for this longer aSD sequence results in an R^2^_adj_ value, relative to the 1^st^ order model, that is larger than previously observed for the smaller 5’-CCUCC-3’ putative aSD sequence (Fig. 3C). Thus, these data indicate that core aSD sequence binding shows a roughly linear relationship with RTE but the inclusion of flanking sequences results in both increasing predictive power as well as increasing non-linearity in the underlying relationship.

**Figure 3:**
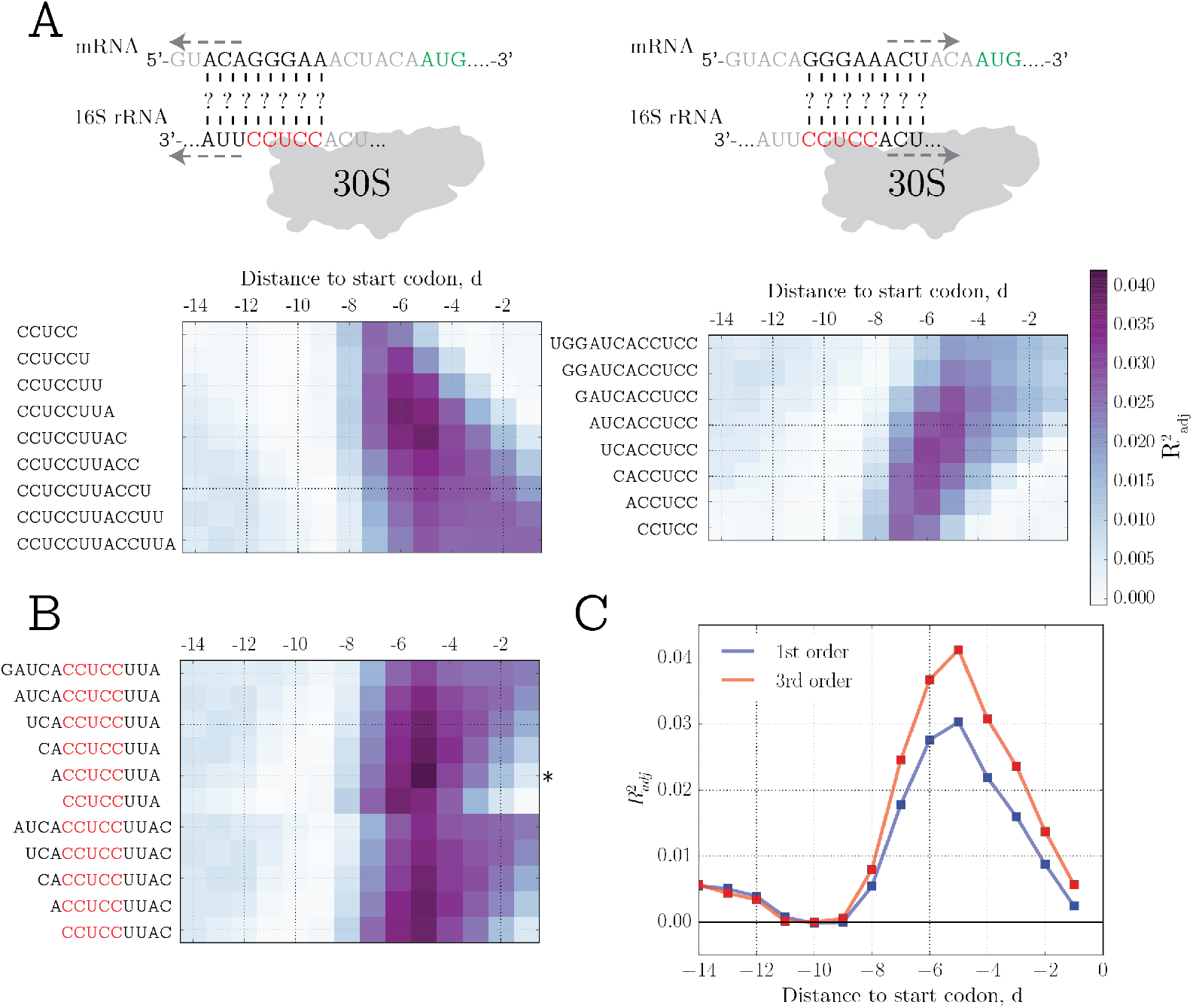
Parameter fitting landscape to determine optimal aSD and distance values. A) R^2^_adj_ from the 3^rd^ order model at different distances to the start codon and various 3’ and 5’ extensions to the core aSD for *E. coli*. B) Combination of best fitting putative aSDs from (A) to determine the optimal aSD sequence and distance parameters based on their fit to the RTE data (* denotes the selected best fitting aSD sequence). C) Comparison of R^2^_adj_ between the 1^st^ and 3^rd^ order polynomial models from the best performing aSD sequence from (B).

### The relationship between aSD binding and translation efficiency

In order to test the generality of our findings for *E. coli*, we next investigated the optimal aSD sequence for two other organisms: *B. subtilis* and *C. crescentus*. We found that the 5’ extensions are similar for the different organisms studied with *B. subtilis* showing preference for a slightly longer 5’ aSD extension, a finding that is in-line with prior observations that the canonical SD sequence in *B. subtilis* 5’ UTRs appears shifted upstream of the start codon (Fig. 1B). We further found that species-specific 3’ extensions to the 16S rRNA continue to result in enhanced correlations and thus are likely to participate in message discrimination for these two organisms (Supporting Figs. 3, 4). For *C. crescentus* the aSD sequence that we obtained from our data-driven model is 5’-CCUCCUUUC-3’ while for *B. subtilis* the corresponding sequence is the extended 5’-UCACCUCCUUUCUA-3’. However, as with *E. coli*, it is difficult to discern whether single base additions/deletions to the ends of these putative aSD sequences are functional (Fig. 4A).

**Figure 4:**
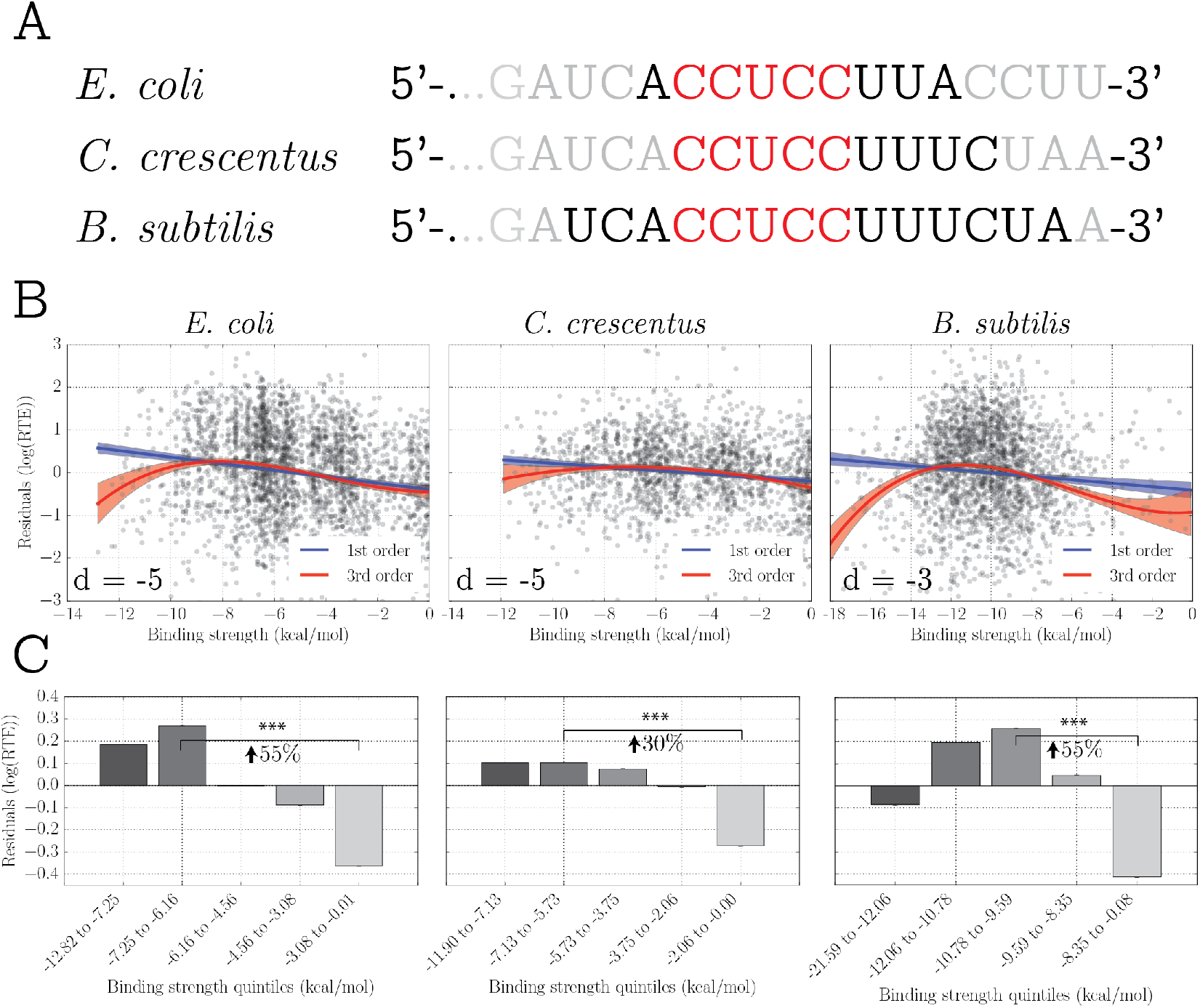
Summary of findings for three independent organisms using ribosome profiling based data. A) Optimal aSD sequences for the three organisms studied. The core sequence is shown in red, and the best fitting aSD is shown in black. B) Best fitting 1^st^ and 3^rd^ order models (blue and red, respectively) for the species-specific optimal aSD sequence and optimal distance to the start codon. 3^rd^ order models produced R^2^_adj_ of 0.041, 0.028, and 0.056 for *E. coli, C. crescentus*, and *B. subtilis* (all cases, p<10^-12^). C) Raw data from B shown averaged within equally sized quintile bins of aSD binding strength. Bars denote the mean within each bin, while error bars show standard error of the mean. Comparison between weakest binding and intermediate-to-strong bins were performed with Wilcoxon rank-sum test (*** denotes *p*<10^-12^), and percent increase highlights the average increase in translation efficiency expected for a gene with a proper strength aSD binding sequence compared to a weakly binding sequence.

Despite the vast evolutionary distance between these species, the functional form of the best fitting models was highly similar for all three, showing the highest values for intermediate binding strengths with similar predictive powers in the 3^rd^ order model (R^2^_adj_ = 0.041, 0.028, and 0.056, for all cases p<10^-12^) (Fig. 4B). We further verified that higher order polynomials provide a superior fit to the data even when penalizing for the increased parameter number via the use of R^2^_adj_ and the Akaike Information Criterion (AIC), a stringent model selection metric used to judge the relative quality of fits (Supporting Fig. 5).

In order to show the magnitude of the observed, we split the data for each organism into equally sized quintile bins (i.e. the 20% strongest binding genes, through to the 20% weakest binding genes). Notably, treating the data this way involves no model fitting and in doing so, we observe that: (i) the average gene which binds the aSD sequence at the intermediate-to-strong binding strength level shows a 30-50% increase in translation efficiency compared to an average gene that binds the aSD very weakly (Fig. 4C) and (ii) the strongest binding quintile of genes exhibits either decreased or equivalent translation efficiency compared to the bin with intermediate-to-strong aSD binding strength. This suggests that sequences that bind too strongly to the aSD sequence may actually show reduced translation efficiency of down stream genes, a point that has support from several prior studies in the literature(Komarova et al., 2002; Ringquist et al., 1995).

Given recent concerns in the literature about the possibility of biases arising from the size selection step of prokaryotic ribosome profiling studies, we analyzed two further *E. coli* datasets (n=1278 and 1321) from an independent lab generated in such a way as to minimize potential sources of error(Mohammad et al., 2016). We observed nearly identical results to the previous *E. coli* dataset (Supporting Fig. 6). For both replicates, the 5’-ACCUCCUUA-3’ aSD sequence at a distance of -5 provided the best fit to the data with corresponding R^2^_ad_j values of 0.06 and 0.07 for the best fitting 3^rd^ order polynomial and effect sizes of 45% and 50%. While illustrating the robustness of our results for a given organism across multiple independent datasets, this analysis also highlights the sensitivity of R^2^_adj_ to noise (we observed a roughly 50% increase with these data) and the relative insensitivity of the effect size.

### Translation efficiency in other data sets

Ribosome profiling provides unprecedented insight into genome-wide translation patterns, but measuring translation efficiency in this manner is error prone. To address this issue, we utilized an independent data set from Taniguchi *et al*. who estimated protein production per mRNA from the green fluorescent protein (GFP)-tagged single-cell protein distributions for 1018 *E. coli* genes (see Materials and Methods) (Taniguchi et al., 2010). Using their data, we performed the same analysis as above and observed nearly identical results to those seen in Fig. 4 for *E. coli*. In other words, the data exhibit a maximum at intermediate-to-strong aSD binding strengths (Fig. 5A,B). When we limit our analysis to genes with progressively more stringent signal to error ratios, the magnitude of the R^2^_adj_ gets larger such that in the 20% of genes with the highest confidence measurements (n=204) we find that 5’-ACCUCCUUA-3’ binding at a spacing of -5 predicts RTE with an R^2^_adj_ of 0.10 (p<10^-5^, compared to R^2^_adj_ =0.064 for the linear fit) (Supporting Fig. 7).

**Figure 5:**
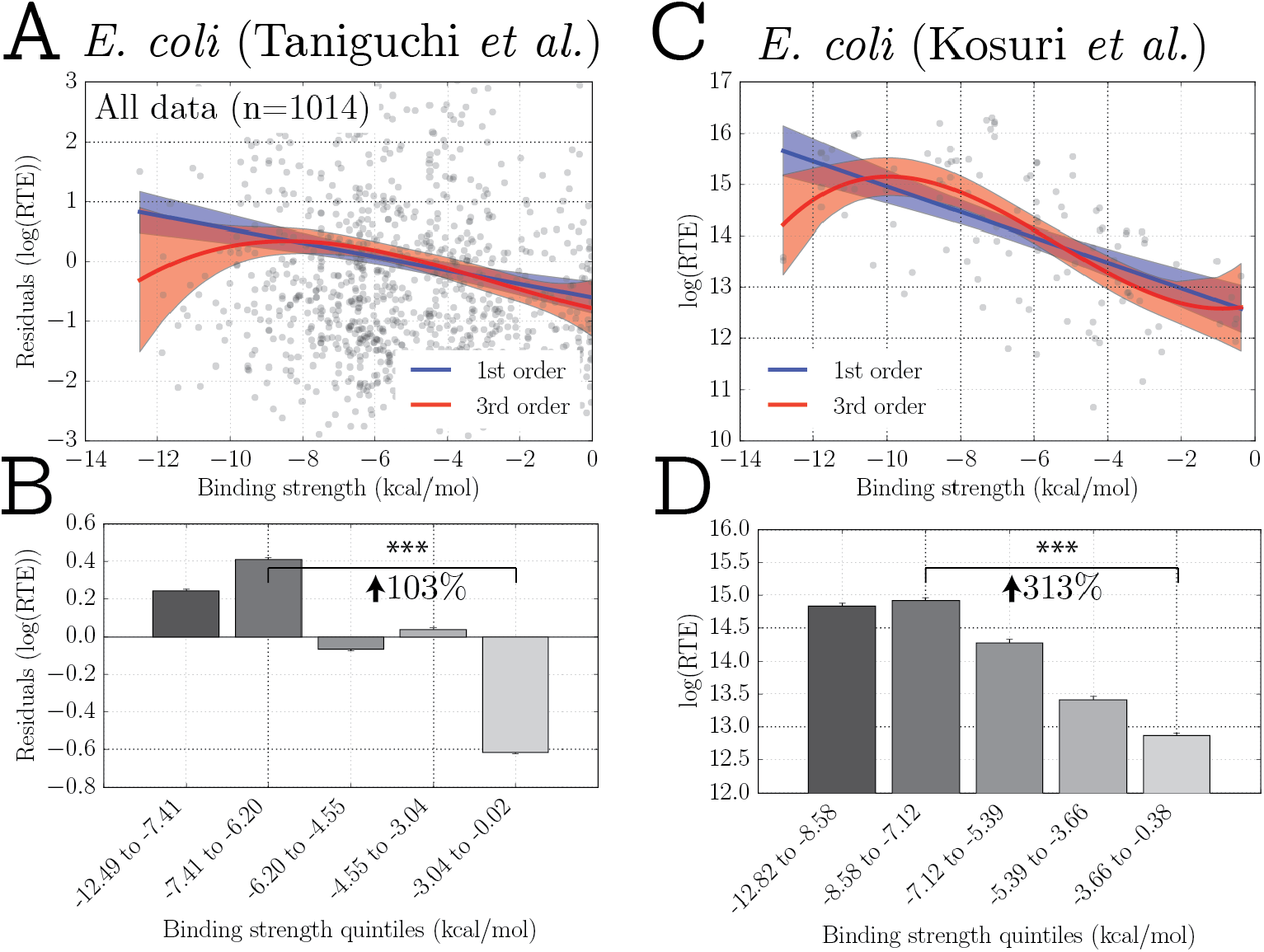
Validation of principal findings in non-ribosomal protein based datasets A) Utilizing the data from Taniguchi *et al*. we observe a significant relationship between aSD binding strength and residual RTE values that mimics ribosome profiling based data (R^2^_adj_ = 0.023 and 0.026 for 1^st^ and 3^rd^ order fits, p<10^−6^ for both cases). B) We also show, using quintile analysis of this data, a 103% increase in RTE between weakly binding and intermediate-to-strong aSD binding genes (*** denotes p<10^-6^, rank-sum test). C) Utilizing recombinant GFP expression data from Kosuri *et al*. we show that a 3^rd^ order model provides a better fit to the data as compared to a 1st order linear model (R^2^_adj_ = 0.32 and 0.37 for 1^st^ and 3^rd^ order models, p<10^-10^ for both cases). D) Quintile analysis of these data shows a large effect size (*** denotes p<10^-6^, rank-sum test) as well as a plateau / slight-decrease for the strongest aSD binding quintile.

Finally, although our interest here is in the relationship between aSD sequence binding to mRNAs of endogenous genes, we further verified our main conclusions using a controlled experimental dataset (Kosuri et al., 2013). Kosuri *et al*. measured the strength of 111 ribosome binding sites (RBS) and 114 promoters by creating constructs whereby each promoter/RBS combination drove expression of a downstream GFP reporter (see Materials and Methods). By measuring the resulting protein and mRNA levels, the authors were able to determine, for each RBS, the average protein per mRNA across the different constructs (as above, we refer to this protein/mRNA metric as RTE for simplicity). We observed that a 3^rd^ order polynomial model provided a better fit to the data than a 1^st^ order linear model (R^2^_adj_ = 0.37 and 0.316, respectively, p<10^−10^ in both cases) (Fig. 5C). We also observed that the intermediate binding quintile produced RTE values 85% higher than the weakest binding quintile and a plateau or slight decrease in RTE for the strongest binding quintile of RBS sequences (Fig. 5D). This provides further support for our conclusion that translation efficiency is maximized at intermediate aSD binding strengths. Although the relationship between aSD binding strength and translation in heterologous genes was previously known, this experimental data confirms our genome-scale findings that having an intermediate strength aSD binding sequence upstream of a gene can significantly enhance downstream translation.

## DISCUSSION

Our work illustrates that there is a strong relation between aSD sequence binding and the translation of endogenous genes. We show that genome-scale methods such as ribosome profiling can be leveraged to refine our understanding of basic mechanisms of translation regulation. Specifically, we demonstrate that (i) aSD binding to mRNA is predictive of translation efficiencies for endogenous genes within a relatively narrow window relative to the start codon, (ii) slight changes in the putative aSD sequence result in an array of different statistical conclusions allowing us to determine a data-driven definition of the optimal aSD for each species and (iii) intermediate binding strengths between the aSD sequence 5’ UTRs maximize the translation efficiency of downstream genes in all datasets that we encountered.

A variety of prior bioinformatics studies have made important contributions to our understanding of translation initiation, however the range of different definitions used to categorize the aSD binding relationship is cause for concern. We show that even small changes in this definition can result in different statistical findings within the same dataset. Indeed, several previous studies failed to show any association between aSD binding strengths and translation efficiency measured via ribosome profiling, a fact that we resolve here and attribute largely to differences in the definition of relevant parameters for this interaction (G.-W. Li et al., 2014; Schrader et al., 2014).

As others have noted, by stabilizing the components necessary for translation initiation, the presence of a strong binding sequence likely prevents premature disassociation of the initiation complex. However, if the aSD sequence binds too tightly to the 5’ UTR, it may be difficult to break this interaction and enter the elongation phase of translation (Li et al., 2012). Our model, which has support from several prior studies in the literature, postulates that the relationship between aSD binding and overall translation efficiency is thus non-linear with translation efficiency maximized at intermediate binding strengths between the aSD sequence and mRNA (Fig. 6) (Guimaraes et al., 2014; Komarova et al., 2002; Ringquist et al., 1995). In each dataset that we analyzed, we observed the same consistent trend; translation efficiencies show a maximum at intermediate aSD binding strengths. Studies that fail to observe this drop-off at high binding strengths may simply fail to access the strongest binding sequences. Since we have shown that the difference between 1^st^ and 3^rd^ order models is relatively minor when investigating binding to short aSD sequences, sequences with perfect complementarity to the core aSD sequence may indeed show a roughly linear increase in translation efficiencies. However, when considering longer aSD sequences perfect complementary becomes detrimental as it begins to include flanking sequences.

**Figure 6:**
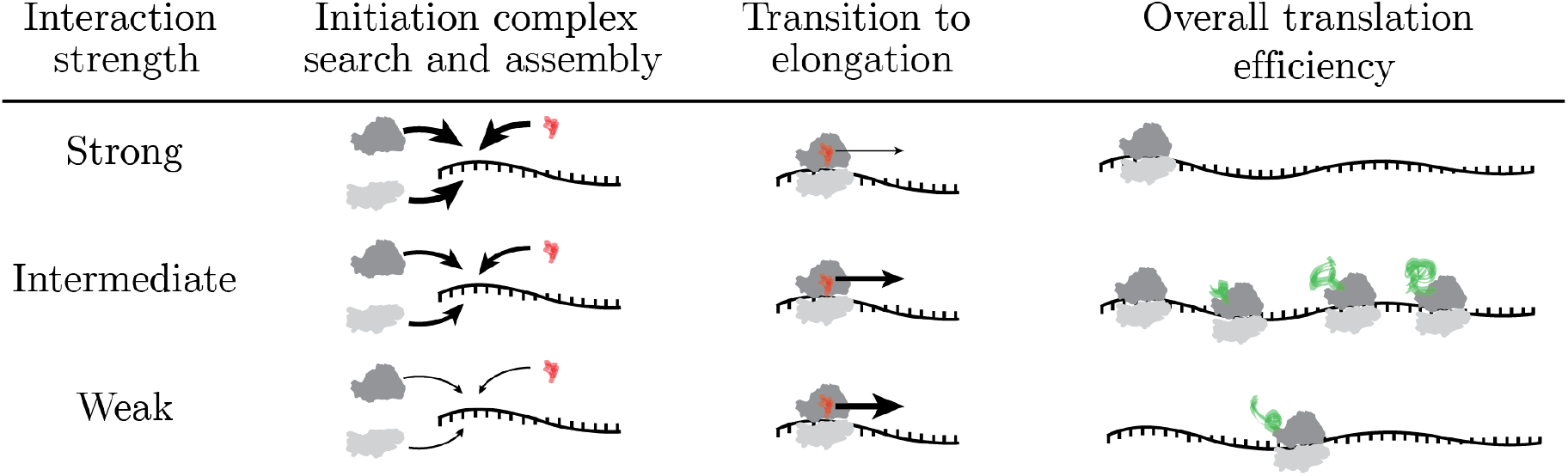
Illustration of our proposed model explaining why translation efficiency is maximized at intermediate levels of aSD binding. The competing processes of initiation complex assembly and transition into elongation select for and against, respectively, strong aSD binding to mRNAs resulting in maximal translation efficiency for intermediate binding strength sequences that balance these processes.

While our methodology allows us to show that there is a significant relationship between aSD binding strength and translation efficiency of downstream genes, it is nevertheless surprising that the predictive power of this relationship is so low given that: (i) the aSD sequence is so highly conserved, and (ii) experimental investigations have seen large changes in protein output when modulating 5’ UTR sequence binding to the aSD via sequence changes in the mRNA(Salis et al., 2009). However, as we have stressed throughout, we note again here that our findings are likely to represent a lower bound on the predictive power of this interaction for several reasons. Genome-scale metrics are subject to both technical and biological noise, and translation efficiency as a metric will particularly suffer from this noise due to error-propagation. Further, mRNA folding around the start codon is known to exert a large affect on translation efficiencies and our computational approximations are at best rough approximations of the true structure for any gene(Park et al., 2013). It is thus reasonable to assume that these sources of noise contribute to lowering the *expected* “perfect” correlations far below 1.0 as has been shown in other systems(Csárdi et al., 2015). Nevertheless, the underlying relationship that we observe is strong enough to overcome these concerns leading to statistically significant correlations in all datasets that we investigated.

Continued development and application of the ribosome profiling technique and associated technologies to diverse organisms will be critical for clarifying a number of outstanding questions in the field of translation regulation. We show that these genome-scale studies can be leveraged to provide insight into basic biological mechanisms and this will only continue as we gain data for more and more species and continue to understand and limit sources of experimental noise. A better understanding of the rules governing translation initiation and translation efficiency from this systems viewpoint has the practical potential to enhance our ability to design and engineer optimal protein expression systems for a host of biotechnological purposes.

## MATERIAL AND METHODS

### The data and relative translation efficiency

We downloaded ribosome profiling reads and corresponding RNA-sequencing reads for *E. coli, C. crescentus, and B. subtilis*(G.-W. Li et al., 2014; Schrader et al., 2014; Subramaniam et al., 2013). For our analysis, we used the original researchers mapping of sequence reads to the respective genomes (.wig files) and removed genes with coverage below 25% in either the RNA-seq or ribosome profiling data-sets in order to enrich for high confidence translation efficiency measurements given the problem of noisy data. We also removed from our analysis any gene shorter than 30 amino acids as well as potentially mis-annotated genes with zero ribosome profiling reads to the first 10 nucleotides.

For all remaining genes that met coverage requirements, we calculated translation efficiency for each gene as the RPKM in the Ribo-seq dataset divided by the RPKM in the RNA-sequencing dataset. As in the text, we emphasize here that our metric of relative translation efficiency derives from dividing two noisy variables and is therefore a rough approximation of the “true” translation efficiency of individual genes. We separately compiled 2 further datasets for *E. coli*, subjecting them to the same pipeline as above(Mohammad et al., 2016). In addition to providing an additional dataset for *E. coli* to test the robustness of the parameters, these data were also collected and mapped in such a way as to purportedly minimize bias associated with long fragment selection and center-weighted mappings, potentially providing a better estimate of the effect size and variance explained by our model.

We further utilize two experimental datasets to independently validate our conclusions. The first from Taniguchi *et al*. utilized single-cell distributions of protein counts to estimate the proteins produced per mRNA from fitted gamma-distributions of single-cell expression. From the original dataset of 1018 genes we remove 4 from our analysis for quality control, i.e. sequences are not a multiple of 3, do not have a ‘product’ annotation’, contain internal stop codons, etc. For clarity we maintain the label of relative translation efficiency (RTE) to describe these data but stress that their derivation is un-related to ribosome profiling based estimates of translation efficiency (Taniguchi et al., 2010) and that RTE in this context has a slightly different interpretation.

We also downloaded detailed experimental data from Kosuri *et al*. who created libraries of Ribosome Binding Site (RBS) and Promoter variants driving Green Fluorescent Protein expression (Kosuri et al., 2013). By measuring mRNA and protein concentrations via FLOW-Seq, each of the 110 RBSs can be described by their protein levels divided by mRNA levels, which are collected for each construct and averaged across all the promoter variants containing this RBS construct (from their initial data we exclude the ‘Dead-RBS’ construct because it's short length is prohibitive to our analysis). Here we analyze this “mean.xlat” data (as described in their supporting tables of Kosuri *et al*.) as a measure of relative translation efficiency. As before, although the ribosome profiling estimates, the data from Taniguchi *et al*. and this data from Kosuri *et al*. are all quantifying translation efficiency in slightly different ways that have slightly different interpretations, each of these metrics should at very least correlate strongly with the idea of translation efficiency as we have defined it so for ease of language we continue to refer to this as relative translation efficiency (RTE). Rather than subtracting out the effect of mRNA structure as in the other studies, we simply provide regressions on this raw data here since a) the downstream gene is the same and thus structure is mostly preserved between constructs and b) each promoter will introduce slightly different sequences upstream of the RBS but their structural effects of this introduction should be accounted for in the averaging process.

### Gene classification and quantification of aSD binding strength

All calculations of RNA folding were performed using the RNAfold method from ViennaRNA with default parameters(Hofacker, 2003). Estimations of cis-structure were based on calculated folding energies for the -30 to +30 nt region relative to the start codon. RNA::RNA hybridizations were performed using the RNAcofold method with default parameters. For each gene, we iterated through all x-mers (where x is the length of the putative aSD sequence) upstream of the start codon in order to capture 14 hybridization events (as this was a reasonable point at which our testing showed essentially no correlations for any aSD sequence as evidenced in Figs. 2 & 3).

### Statistics and code sharing

All code used to perform translation efficiency measurements, as well as all statistics were written using custom scripts in Python that are freely available. All regression models (including R^2^_ad_j and Akaike Information Criteria (AIC)) were based off fits using the statsmodels package; reported p-values in all regressions are from the F-test.

## AUTHOR CONTRIBUTIONS

AJH, LANA, and MCJ conceived the project. AJH designed and performed the experiments, analyzed the results, prepared the figures and wrote the manuscript. ARP, LANA and MCJ supervised the research, provided critical feedback and made substantial edits to the manuscript.

## ACKNOWLEDGMENTS

The authors thank Peter Winter, João Moreira, and Sophia Liu for critical reading of the manuscript.

## Supporting Information

**Figure S1:**
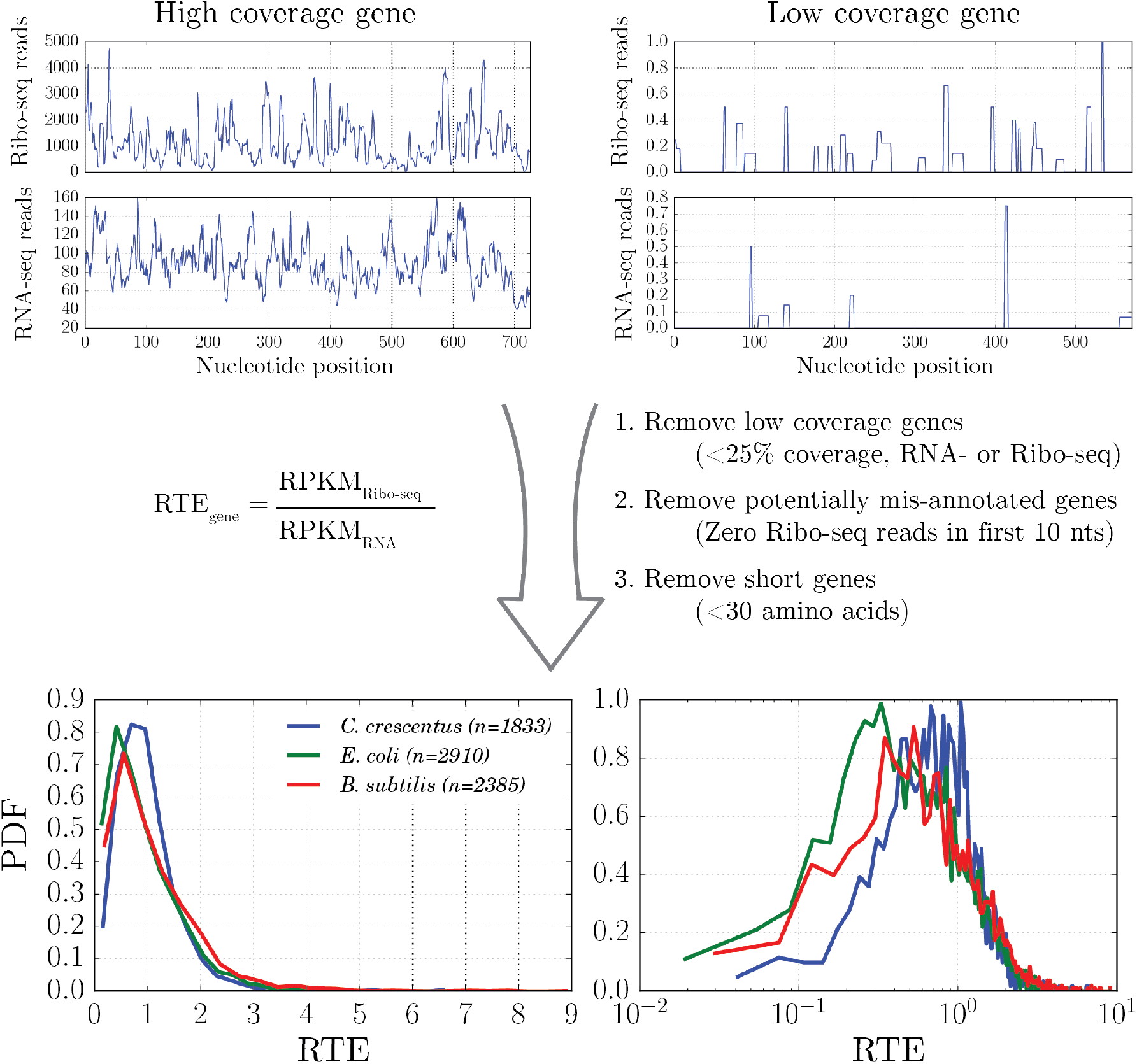
Example gene profiles showing mapped RNA- and Ribo-seq reads that are used as input to calculate RTE. Our pipeline first removes a subset of the total genes based off of coverage, annotation, and length requirements resulting in RTE measurements for 2910, 1833, and 2385 genes in *E. coli, C. crescentus* and *B. subtilis*. Distributions of the RTE values on normal and log-scale show that RTE is approximately log-normally distributed and comparable between the three datasets studied.

**Figure S2:**
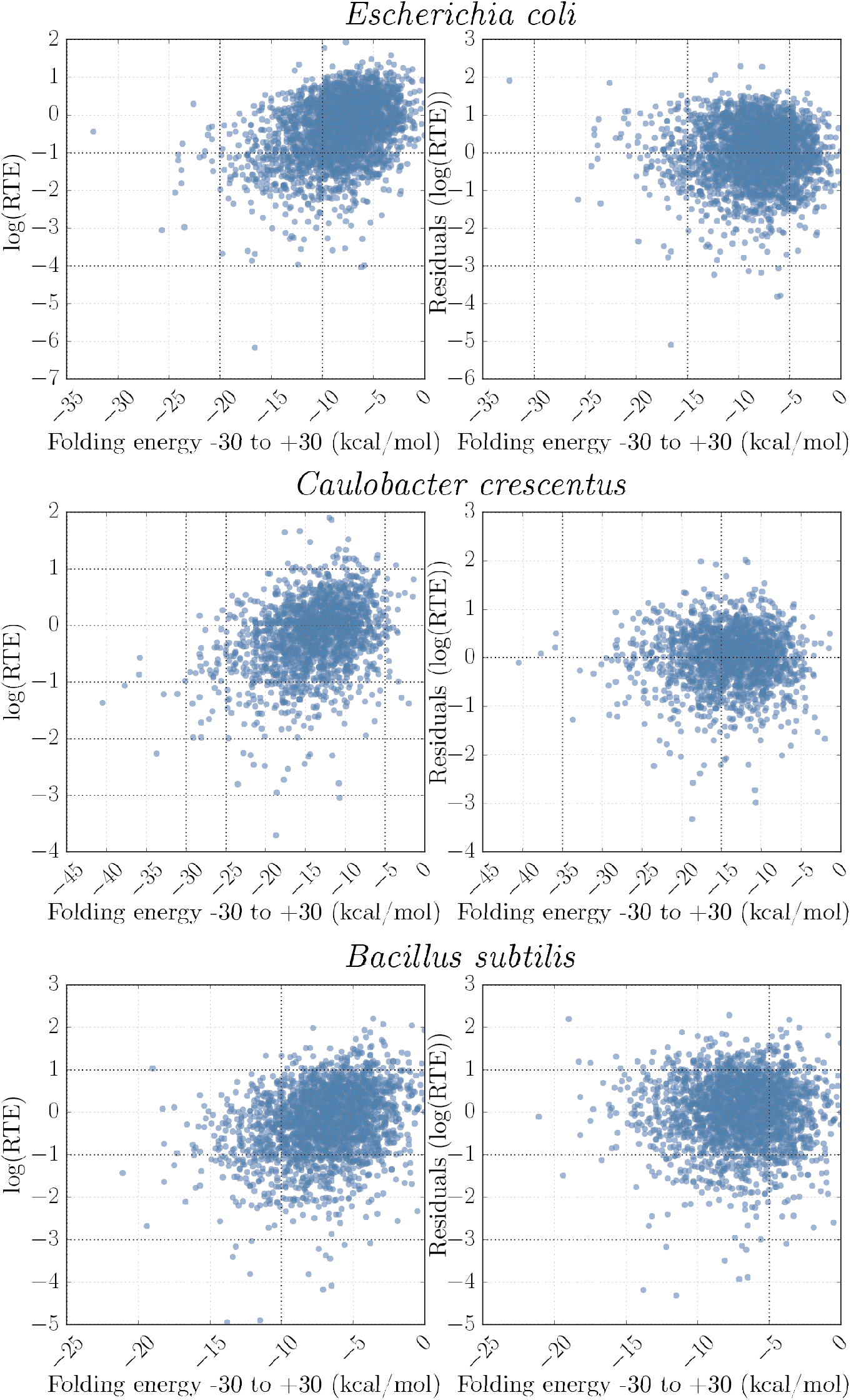
Correlation between the free energy of RNA folding around the start codon (-30 to +30) and log(RTE) for three different organisms studied (left R^2^ =0.13, 0.10, 0.08 for *E. coli, C. crescentus*, and *B. subtilis* respectively; for all cases p<10^-43^). For RTE in the main text we utilize the residuals from the best fitting linear model based off this regression for each organism, effectively removing the influence of mRNA structure on RTE (right).

**Figure S3; related to Fig. 3:**
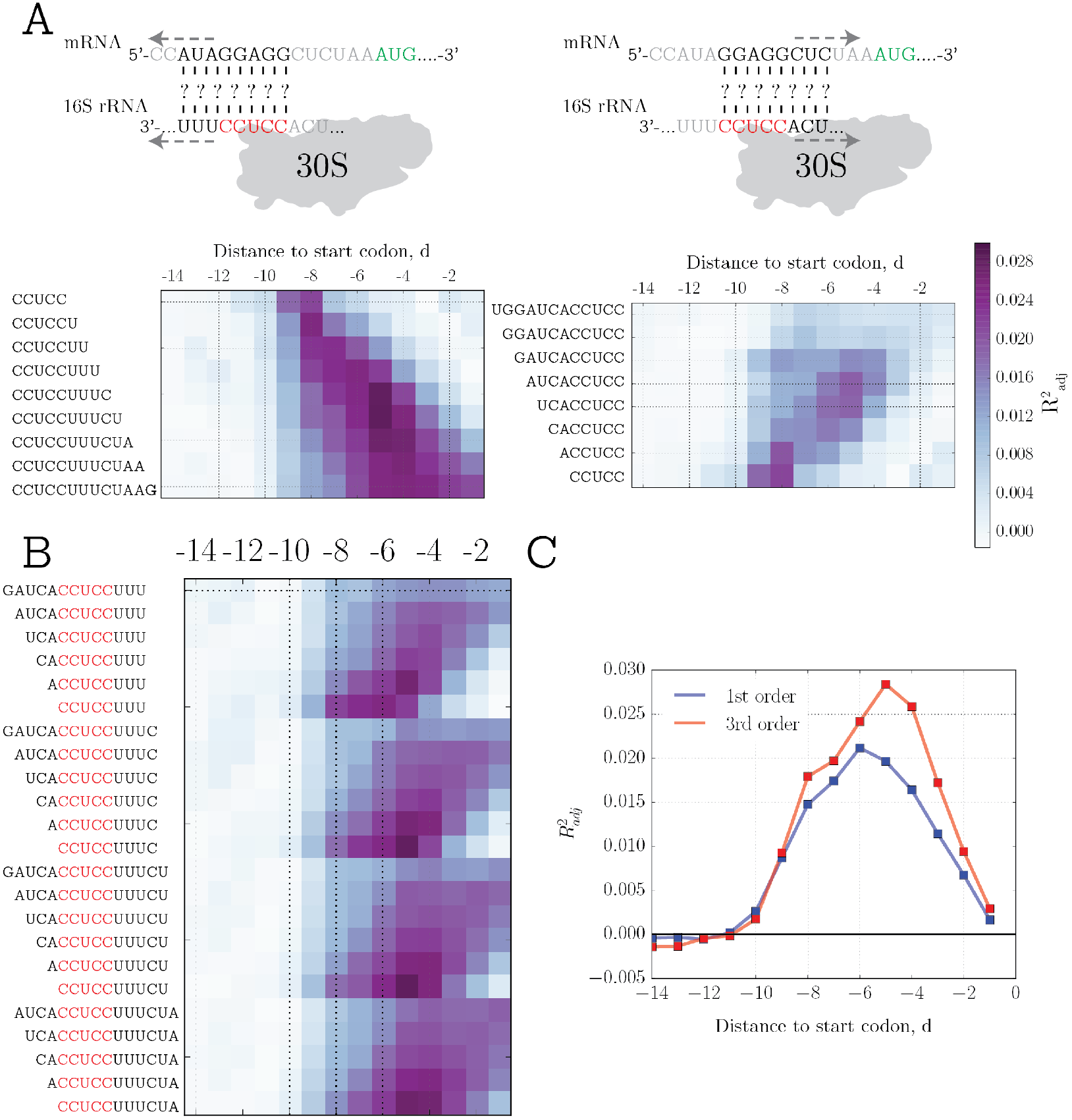
A) R^2^_adj_ from the 3^rd^ order model at different distances to the start codon and various 3, and 5, extensions to the core aSD for *C. crescentus*. B) Combination of best fitting putative aSDs from (A) to determine the optimal aSD sequence. C) Comparison of R^2^_adj_ between the 1^st^ and 3^rd^ order polynomial models from best performing aSD sequence.

**Figure S4; related to Figure 3:**
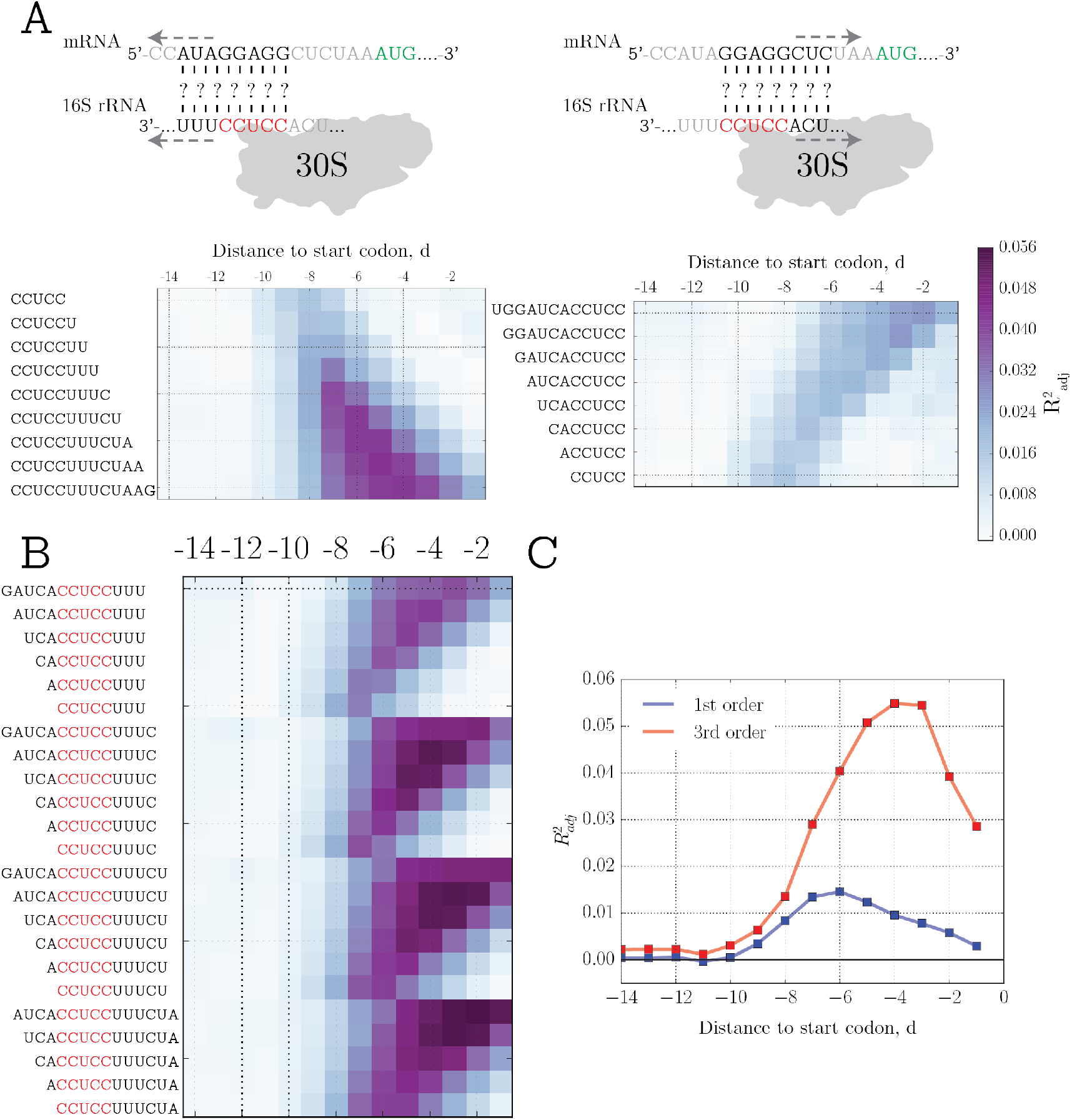
A) R^2^_adj_ from the 3^rd^ order model at different distances to the start codon and various 3, and 5, extensions to the core aSD for *B. subtilis*. B) Combination of best fitting putative aSDs from (A) to determine the optimal aSD sequence. C) Comparison of R^2^_adj_ between the 1^st^ and 3^rd^ order polynomial models from best performing aSD sequence.

**Figure S5; related to Figure 4:**
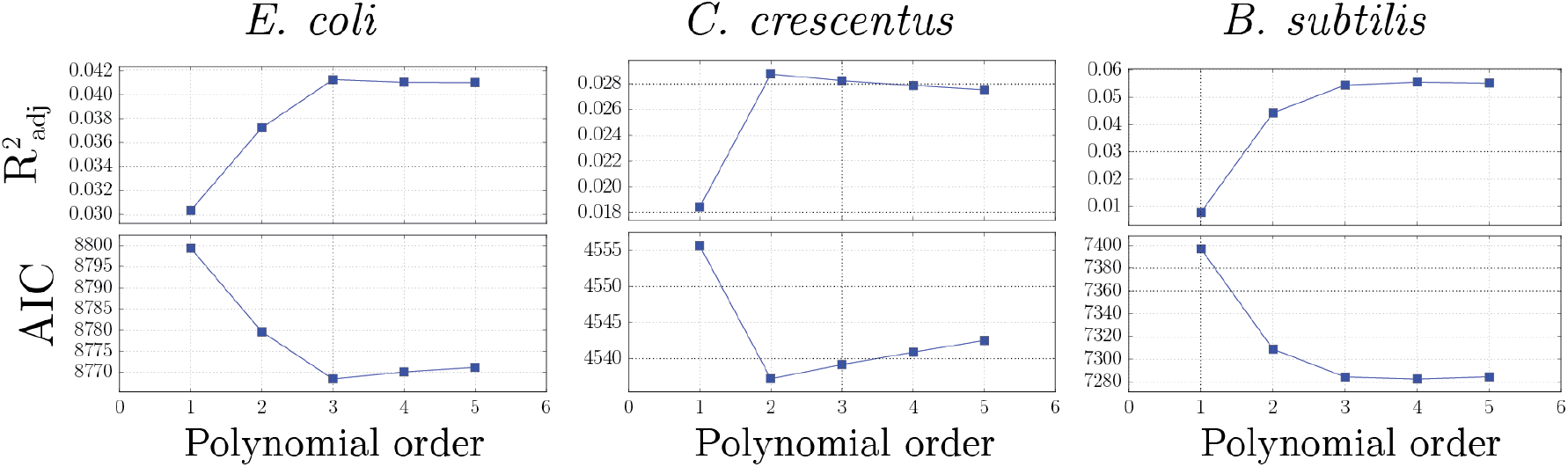
For *E. coli* data (left), given the optimal distance and aSD parameters, we show the effect of increasingly complex polynomial fits on the R^2^_adj_ (top) and Akaike Information Criterion (AIC)(bottom), two statistical methods commonly used for model selection. Both metrics penalize models with increasing parameter number through different statistical means in order to prevent over-fitting; the best model, according to the R^2adj^, should be the one that maximizes this metric while for the AIC the best model should minimize this value. The data are also shown for *C. crescentus* (center) and *B. subtilis* (right) data, in all cases this data was calculated using the optimal aSD and spacing values indicated in Fig. 4 of the main text for each organism.

**Figure S6:**
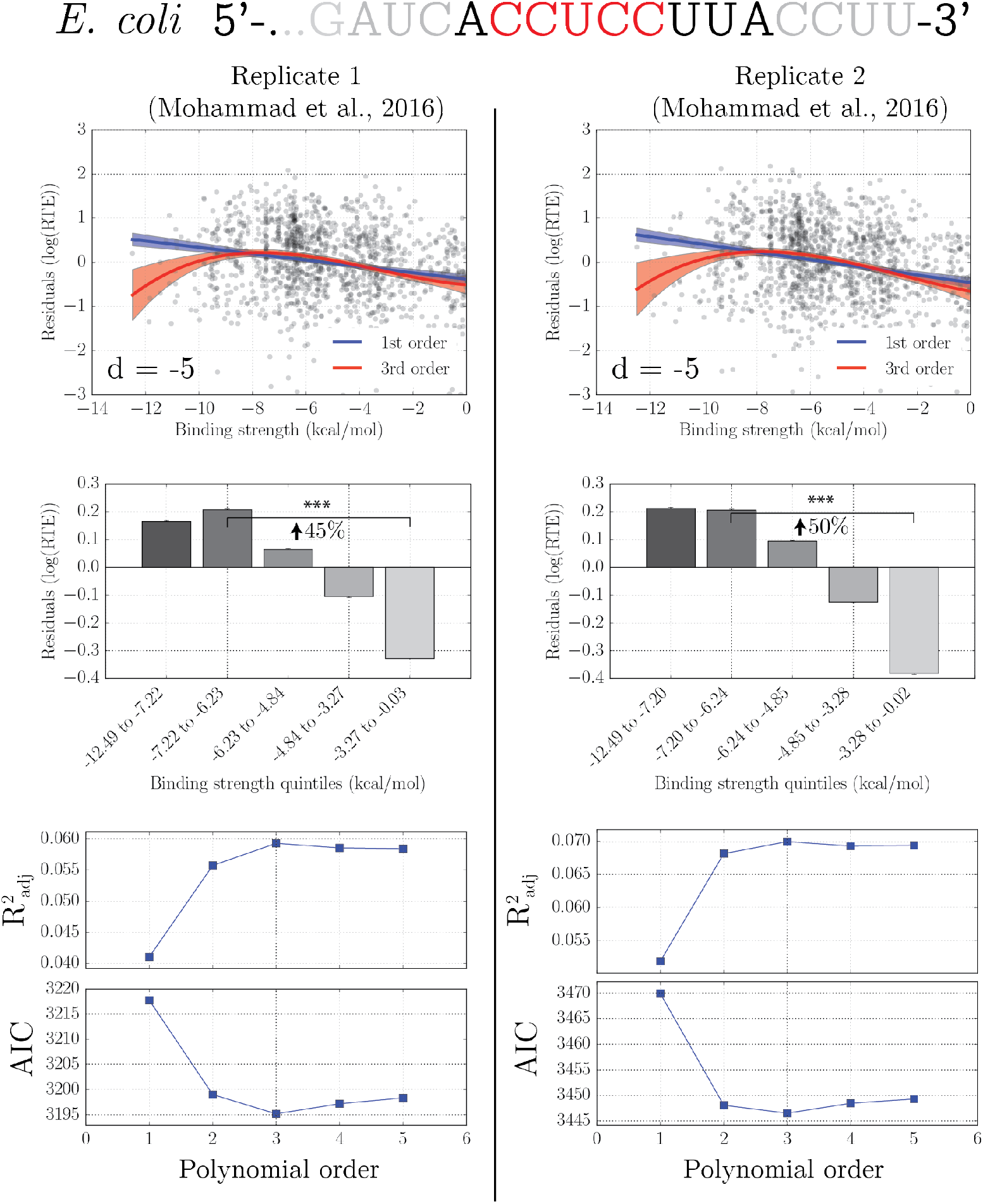
Replication of *E. coli* findings in independent datasets from Mohammad *et al*. We repeated the process shown in Fig. 3 of the main text and for both datasets and determined that the optimal aSD parameters and distance corresponded to 5,-ACCUCCUUA-3, at a distance of -5, as was found in Fig. 3 for *E. coli*. Top, as in Fig. 4 of the main text, we show fits to the residual RTE values (R^2^_adj_ for 3^rd^ order fits = 0.06 and 0.07, respectively). Middle, as in Fig. 4 of the main text we show the quintile analysis with respective effect sizes for equally sized bins. Bottom, as in Supporting Fig. S5 we show the relevant model selection parameters for determining the best fitting model.

**Figure S7; related to Figure 5:**
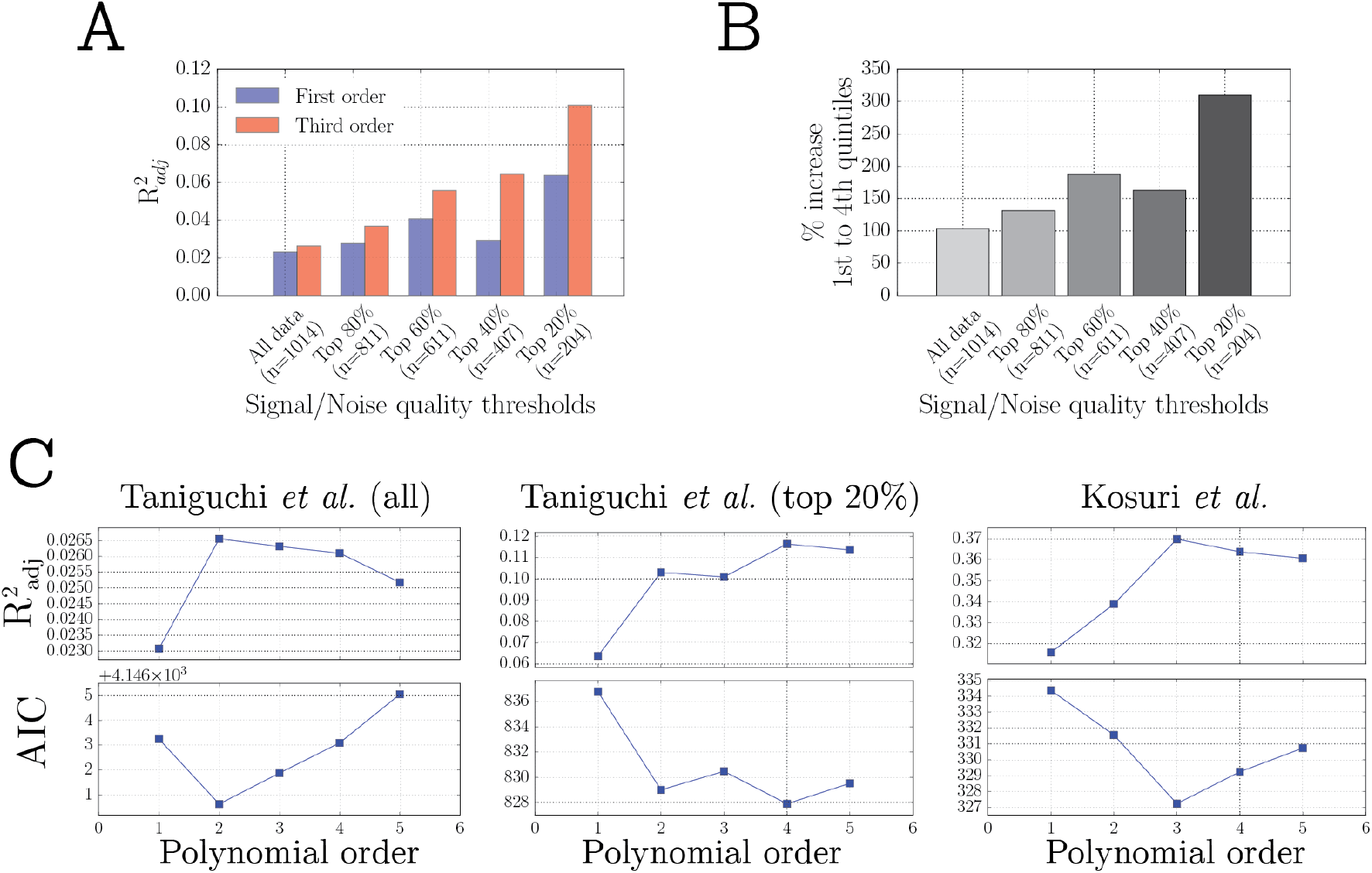
A) Restricting the dataset from Taniguchi *et al*. based on signal to noise thresholds, we observe increasingly better model fits for higher quality data. B) The magnitude of the effect between weak and intermediate-to-strong aSD binding sequences also increases for higher quality data. C) As in Supporting Fig. S5 we show the relevant model selection parameters for determining the best fitting model for the three-labeled datasets.

## REFERENCES

Barendt, P.A., Shah, N.A., Barendt, G.A., Kothari, P.A., Sarkar, C.A., 2013. Evidence for context-dependent complementarity of non-shine-dalgarno ribosome binding sites to Escherichia coli rRNA. ACS Chem. Biol. 8, 958–966. doi:10.1021/cb3005726

Barrick, D., Villanueba, K., Childs, J., Kalil, R., Schneider, T.D., Lawrence, C.E., Gold, L., Stormo, G.D., 1994. Quantitative analysis of ribosome binding sites in E.coli. Nucleic Acids Res. 22, 1287–95.

Bentele, K., Saffert, P., Rauscher, R., Ignatova, Z., Blüthgen, N., 2013. Efficient translation initiation dictates codon usage at gene start. Mol. Syst. Biol. 9, 1–10. doi:10.1038/msb.2013.32

Calogero, R.A., Pon, C.L., Canonaco, M.A., Gualerzi, C.O., 1988. Selection of the mRNA translation initiation region by Escherichia coli ribosomes. Proc. Natl. Acad. Sci. U. S. A. 85, 6427–6431. doi:10.1073/pnas.85.17.6427

Chang, B., Halgamuge, S., Tang, S.L., 2006. Analysis of SD sequences in completed microbial genomes: Non-SD-led genes are as common as SD-led genes. Gene 373, 90–99. doi:10.1016/j.gene.2006.01.033

Chen, H., Bjerknes, M., Kumar, R., Jay, E., 1994. Determination of the optimal aligned spacing between the Shine-Dalgarno sequence and the translation initiation codon of Escherichia coli mRNAs. Nucleic Acids Res. 22, 4953–4957.

Cortes, T., Schubert, O.T., Rose, G., Arnvig, K.B., Comas, I., Aebersold, R., Young, D.B., 2013. Genome-wide mapping of transcriptional start sites defines an extensive leaderless transcriptome in Mycobacterium tuberculosis. Cell Rep. 5, 1121–31. doi:10.1016/j.celrep.2013.10.031

Csárdi, G., Franks, A., Choi, D.S., Airoldi, E.M., Drummond, D.A., 2015. Accounting for experimental noise reveals that mRNA levels, amplified by post-transcriptional processes, largely determine steady-state protein levels in yeast. PLOS Genet. 11, e1005206. doi:10.1371/journal.pgen.1005206

de Smit, M.H., van Duin, J., 1994. Translation initiation on structured messengers: another role for the Shine-Dalgarno interaction. J. Mol. Biol. 235, 173–184.

de Smit, M.H., van Duin, J., 1990. Secondary structure of the ribosome binding site determines translational efficiency : A quantitative analysis. Proc. Natl. Acad. Sci. 87, 7668–7672.

Dekel, E., Alon, U., 2005. Optimality and evolutionary tuning of the expression level of a protein. Nature 436, 588–92. doi:10.1038/nature03842

Espah Borujeni, A., Channarasappa, A.S., Salis, H.M., 2014. Translation rate is controlled by coupled trade-offs between site accessibility, selective RNA unfolding and sliding at upstream standby sites. Nucleic Acids Res. 42, 2646–59. doi:10.1093/nar/gkt1139

Goodman, D.B., Church, G.M., Kosuri, S., 2013. Causes and effects of N-terminal codon bias in bacterial genes. Science (80-.). 342, 475–479.

Gu, W., Zhou, T., Wilke, C.O., 2010. A universal trend of reduced mRNA stability near the translation-initiation site in prokaryotes and eukaryotes. PLoS Comput. Biol. 6, e1000664. doi:10.1371/journal.pcbi.1000664

Guimaraes, J.C., Rocha, M., Arkin, A.P., 2014. Transcript level and sequence determinants of protein abundance and noise in Escherichia coli. Nucleic Acids Res. 42, 4791–9. doi:10.1093/nar/gku126

Hockenberry, A.J., Sirer, M.I., Amaral, L.A.N., Jewett, M.C., 2014. Quantifying position-dependent codon usage bias. Mol. Biol. Evol. 31, 1880–93. doi:10.1093/molbev/msu126

Hofacker, I.L., 2003. Vienna RNA secondary structure server. Nucleic Acids Res. 31, 3429–3431. doi:10.1093/nar/gkg599

Ingolia, N.T., Ghaemmaghami, S., Newman, J.R.S., Weissman, J.S., 2009. Genome-wide analysis in vivo of translation with nucleotide resolution using ribosome profiling. Science 324, 218–23. doi:10.1126/science.1168978

Komarova, A. V, Tchufistova, L.S., Supina, E. V, Boni, I. V, 2002. Protein S1 counteracts the inhibitory effect of the extended Shine-Dalgarno sequence on translation. RNA 8, 1137–1147. doi:10.1017/S1355838202029990

Kosuri, S., Goodman, D.B., Cambray, G., Mutalik, V.K., Gao, Y., Arkin, A.P., Endy, D., Church, G.M., 2013. Composability of regulatory sequences controlling transcription and translation in Escherichia coli. Proc. Natl. Acad. Sci. 110, 14024–9. doi:10.1073/pnas.1301301110

Kramer, P., Gäbel, K., Pfeiffer, F., Soppa, J., 2014. Haloferax volcanii, a prokaryotic species that does not use the Shine Dalgarno mechanism for translation initiation at 5’-UTRs. PLoS One 9, e94979. doi:10.1371/journal.pone.0094979

Kudla, G., Murray, A.W., Tollervey, D., Plotkin, J.B., 2009. Coding-sequence determinants of gene expression in Escherichia coli. Science 324, 255–258.

Lahens, N.F., Kavakli, I.H., Zhang, R., Hayer, K., Black, M.B., Dueck, H., Pizarro, A., Kim, J., Irizarry, R., Thomas, R.S., Grant, G.R., Hogenesch, J.B., 2014. IVT-seq reveals extreme bias in RNA-sequencing. Genome Biol. 15, R86. doi:10.1186/gb-2014-15-6-r86

Li, G.-W., 2015. How do bacteria tune translation efficiency? Curr. Opin. Microbiol. 24, 66–71. doi:10.1016/j.mib.2015.01.001

Li, G.-W., Burkhardt, D., Gross, C., Weissman, J.S., 2014. Quantifying absolute protein synthesis rates reveals principles underlying allocation of cellular resources. Cell 157, 624–35. doi:10.1016/j.cell.2014.02.033

Li, G.-W., Oh, E., Weissman, J.S., 2012. The anti-Shine–Dalgarno sequence drives translational pausing and codon choice in bacteria. Nature 484, 538–541. doi:10.1038/nature10965

Li, J.J., Bickel, P.J., Biggin, M.D., 2014. System wide analyses have underestimated protein abundances and the importance of transcription in mammals. PeerJ 2, e270. doi:10.7717/peerj.270

Lim, K., Furuta, Y., Kobayashi, I., 2012. Large variations in bacterial ribosomal RNA genes. Mol. Biol. Evol. 29, 2937–2948. doi:10.1093/molbev/mss101

Lu, P., Vogel, C., Wang, R., Yao, X., Marcotte, E.M., 2007. Absolute protein expression profiling estimates the relative contributions of transcriptional and translational regulation. Nat. Biotechnol. 25, 117–24. doi:10.1038/nbt1270

Ma, J., Campbell, A., Karlin, S., 2002. Correlations between Shine-Dalgarno sequences and gene features such as predicted expression levels and operon structures. J. Bacteriol. 184, 5733–5745. doi:10.1128/JB.184.20.5733-5745.2002

Miettinen, T.P., Bjorklund, M., 2014. Modified ribosome profiling reveals high abundance of ribosome protected mRNA fragments derived from 3’ untranslated regions. Nucleic Acids Res. 43, 1019–1034. doi:10.1093/nar/gku1310

Mohammad, F., Woolstenhulme, C.J., Green, R., Buskirk, A.R., 2016. Clarifying the Translational Pausing Landscape in Bacteria by Ribosome Profiling. Cell Rep. 1–9. doi:10.1016/j.celrep.2015.12.073

Mutalik, V.K., Guimaraes, J.C., Cambray, G., Mai, Q.-A., Christoffersen, M.J., Martin, L., Yu, A., Lam, C., Rodriguez, C., Bennett, G., Keasling, J.D., Endy, D., Arkin, A.P., 2013. Quantitative estimation of activity and quality for collections of functional genetic elements. Nat. Methods 10, 347–53. doi:10.1038/nmeth.2403

Na, D., Lee, S., Lee, D., 2010. Mathematical modeling of translation initiation for the estimation of its efficiency to computationally design mRNA sequences with desired expression levels in prokaryotes. BMC Syst. Biol. 4, 71. doi:10.1186/1752-0509-4-71

Nakagawa, S., Niimura, Y., Miura, K., Gojobori, T., 2010. Dynamic evolution of translation initiation mechanisms in prokaryotes. Proc. Natl. Acad. Sci. 107, 6382–6387. doi:10.1073/pnas.1002036107

Park, C., Chen, X., Yang, J.-R., Zhang, J., 2013. Differential requirements for mRNA folding partially explain why highly expressed proteins evolve slowly. Proc. Natl. Acad. Sci. U. S. A. 110, E678–86. doi:10.1073/pnas.1218066110

Ringquist, S., Jones, T., Snyder, E.E., Gibson, T., Boni, I., Gold, L., 1995. High-affinity RNA ligands to Escherichia coli ribosomes and ribosomal protein S1: comparison of natural and unnatural binding sites. Biochemistry 34, 3640–3648.

Rinke-Appel, J., Junke, N., Brimacombe, R., Lavrik, I., Dokudovskaya, S., Dontsova, O., Bogdanov, A., 1994. Contacts between 16S ribosomal RNA and mRNA, within the spacer region separating the AUG initiator codon cross-linking study. Nucleic Acids Res. 22, 3018–3025.

Sakai, H., Imamura, C., Osada, Y., Saito, R., Washio, T., Tomita, M., 2001. Correlation between Shine-Dalgarno sequence conservation and codon usage of bacterial genes. J. Mol. Evol. 52, 164–170. doi:10.1007/s002390010145

Salis, H.M., Mirsky, E.A., Voigt, C.A., 2009. Automated design of synthetic ribosome binding sites to control protein expression. Nat. Biotechnol. 27, 946–50. doi:10.1038/nbt.1568

Schrader, J.M., Zhou, B., Li, G.-W., Lasker, K., Childers, W.S., Williams, B., Long, T., Crosson, S., McAdams, H.H., Weissman, J.S., Shapiro, L., 2014. The coding and noncoding architecture of the Caulobacter crescentus genome. PLoS Genet. 10, e1004463. doi:10.1371/journal.pgen.1004463

Schwanhäusser, B., Busse, D., Li, N., Dittmar, G., Schuchhardt, J., Wolf, J., Chen, W., Selbach, M., 2011. Global quantification of mammalian gene expression control. Nature 473, 337–342. doi:10.1038/nature10098

Shine, J., Dalgarno, L., 1974. The 3’-terminal sequence of Escherichia coli 16S ribosomal RNA: Complementarity to nonsense triplets and ribosome binding sites. Proc. Natl. Acad. Sci. 71, 1342–1346.

Starmer, J., Stomp, A., Vouk, M., Bitzer, D., 2006. Predicting Shine-Dalgarno sequence locations exposes genome annotation errors. PLoS Comput. Biol. 2, 454–466. doi:10.1371/journal.pcbi.0020057

Steijger, T., Abril, J.F., Engström, P.G., Kokocinski, F., Akerman, M., Alioto, T., Ambrosini, G., Antonarakis, S.E., Behr, J., Bertone, P., 2013. Assessment of transcript reconstruction methods for RNA-seq. Nat. Methods 10, 1177–84. doi:10.1038/nmeth.2714

Subramaniam, A.R., DeLoughery, A., Bradshaw, N., Chen, Y., O’Shea, E., Losick, R., Chai, Y., 2013. A serine sensor for multicellularity in a bacterium. Elife 2, 1–17. doi:10.7554/eLife.01501

Taniguchi, Y., Choi, P.J., Li, G.-W., Chen, H., Babu, M., Hearn, J., Emili, A., Xie, X.S., 2010. Quantifying E. coli proteome and transcriptome with single-molecule sensitivity in single cells. Science 329, 533–8. doi:10.1126/science.1188308

Vogel, C., de Sousa Abreu, R., Ko, D., Le, S.-Y., Shapiro, B.A., Burns, S.C., Sandhu, D., Boutz, D.R., Marcotte, E.M., Penalva, L.O., 2010. Sequence signatures and mRNA concentration can explain two-thirds of protein abundance variation in a human cell line. Mol. Syst. Biol. 6, 1–9. doi:10.1038/msb.2010.59

Vogel, C., Marcotte, E.M., 2012. Insights into the regulation of protein abundance from proteomic and transcriptomic analyses. Nat. Rev. Genet. 13, 227–232. doi:10.1038/nrg3185

Zheng, X., Hu, G.-Q., She, Z.-S., Zhu, H., 2011. Leaderless genes in bacteria: clue to the evolution of translation initiation mechanisms in prokaryotes. BMC Genomics 12, 361. doi:10.1186/1471-2164-12-361

Zupanic, A., Meplan, C., Grellscheid, S.N., Mathers, J.C., Kirkwood, T.B.L., Hesketh, J.E., Shanley, D.P., 2014. Detecting translational regulation by change point analysis of ribosome profiling data sets. RNA 20, 1507–18. doi:10.1261/rna.045286.114

